# VirB, a key transcriptional regulator of virulence plasmid genes in *Shigella flexneri*, forms DNA-binding site dependent foci in the bacterial cytoplasm

**DOI:** 10.1101/2020.11.20.392365

**Authors:** Jillian N. Socea, Grant R. Bowman, Helen J. Wing

## Abstract

VirB is a key regulator of virulence genes located on the large virulence plasmid (pINV) of the bacterial pathogen *Shigella flexneri.* VirB is unusual in that it is not related to other transcriptional regulators, instead, it belongs to a protein family that primarily functions in plasmid and chromosome partitioning; exemplified by ParB. Despite this, VirB does not function to segregate DNA, but rather counters transcriptional silencing of virulence genes mediated by the nucleoid structuring protein, H-NS. Since ParB localizes subcellularly as discrete foci in the bacterial cytoplasm, we chose to investigate the subcellular localization of VirB to gain novel insight into how VirB functions as a transcriptional anti-silencer. To do this, a GFP-VirB fusion that retains the regulatory activity of VirB and yet, does not undergo significant protein degradation in S. *flexneri,* was used. Surprisingly, discrete fluorescent foci were observed in live wild-type S. *flexneri* cells and an isogenic *virB* mutant using fluorescence microscopy. In contrast, foci were rarely observed (<10%) in cells cured of pINV. Moreover, in the context of the fusion, amino acid substitutions in the DNA binding domain of VirB resulted in the fluorescent signal becoming entirely diffuse. Combined, these data demonstrate that the VirB:DNA interactions required for the transcriptional anti-silencing activity of VirB on pINV are a prerequisite for the subcellular localization of VirB in the bacterial cytoplasm. The significance of these findings, in light of the anti-silencing activity of VirB, is discussed.

**Importance:** This study reveals the subcellular localization of VirB, a key transcriptional regulator of virulence genes found on the large virulence plasmid in *Shigella.* Fluorescent signals generated by an active GFP-VirB fusion form 2, 3, or 4 discrete foci in the bacterial cytoplasm, predominantly at the quarter cell position. These signals are completely dependent upon VirB interacting with its DNA binding site found either on the virulence plasmid or an engineered surrogate. Our findings: 1) provide novel insight into VirB:pINV interactions, 2) suggest that VirB may have utility as a DNA marker, and 3) raise questions about how and why this anti-silencing protein that controls virulence gene expression on pINV of *Shigella* spp. forms discrete foci/hubs within the bacterial cytoplasm.

## Introduction

Bacterial cells were once viewed as tiny compartments containing mixtures of macromolecules without organization or arrangement. With advances in the creation and use of fluorescent fusion proteins, the discrete subcellular location of many proteins has been revealed (1–6). Such studies have frequently shown that rather than being confined to a particular subcellular address or target, protein positioning may be dynamic, changing throughout the cell cycle (7–11). As such, subcellular localization studies, when properly controlled, can provide a more holistic view of the molecular biology that occurs within cells and generate new insight into protein function and their interactions with other macromolecules.

In *Shigella spp.,* the causative agents of bacillary dysentery (12), the transcriptional regulator VirB is produced in response to human body temperature (13, 14). At 37°C, VirB upregulates the expression of about 50 genes on the large virulence plasmid, pINV (similar effects are seen if *virB* is artificially induced at lower temperatures; (15)). These genes include those encoding the type three secretion system (i.e., the needle and initial effectors), other virulence-associated factors (i.e., IcsP, OspZ, OspD1; (16–18)), and the transcriptional activator MxiE and its coactivator IpgC (15). Perhaps not surprisingly, VirB is essential for the virulence of this important group of pathogens (13, 19).

As an anti-silencing protein, VirB does not function as a canonical transcription factor, but rather functions to offset transcriptional silencing mediated by the histone-like nucleoid structuring protein, H-NS, on pINV (16, 20–23). H-NS binds, coats and condenses AT-rich, horizontally acquired DNA, in a process that has been coined xenogeneic silencing (24–26). Site-specific VirB-DNA interactions likely remodel H-NS:DNA complexes via a poorly understood mechanism, allowing inaccessible DNA or transcriptionally non-permissive DNA to be accessed/engaged by RNA polymerase so that transcription can occur (27). As such, the bacterial processes of transcriptional silencing and anti-silencing are reminiscent of chromatin remodeling in eukaryotic cells. In recent years, it has become clear that transcriptional silencing and anti-silencing is widespread in bacteria (28) and that these opposing regulatory processes are necessary for bacterial fitness because they control many aspects of bacterial cell physiology, including virulence in a variety of pathogens (28–34).

Our recent work has focused on gaining mechanistic insight into transcriptional anti-silencing by VirB (16, 18, 23, 35, 36). To aid these studies, we have looked to proteins that regulate transcription like VirB, but also to proteins that are paralogues of VirB. While classical transcription factors bind to specific DNA motifs to regulate the expression of target genes, how they reach their sites can vary. Some transcription factors are thought to diffuse in 3D space, while others slide along DNA or undergo intersegmental hopping to reach their cognate sites (37). Regardless, the paradigm has been that these proteins are diffuse while searching for their sites and, once they find their sites, which are frequently scattered around the genome, are at concentrations too low to be detected by standard fluorescence microscopy. In contrast, some DNA-binding proteins, including members of the ParB/Spo0J protein superfamily, to which VirB belongs, show a discrete subcellular localization pattern in bacteria during plasmid and/or chromosome partitioning (38–41). These proteins bind to their DNA recognition sites, form multimeric complexes and then, through their association with the ParA ATPase, move to the poles with the DNA prior to cell division; an activity observed in *Corynebacterium glutamicum* and *Bacillus subtilis* (42–45).

In this study, we chose to examine the subcellular localization of VirB. Our goal was to broaden our understanding of VirB:DNA interactions on a cellular level so that we might consider how these interactions underpin its anti-silencing activity. We were intrigued to know if the signals associated with VirB would resemble those expected for a transcription factor or members of the ParB superfamily. If a subcellular localization signal was detected, we reasoned that this study would become foundational for future work that addresses which attributes of VirB are needed for this localization, whether other macromolecules are required, and ultimately how these factors affect the regulation of virulence genes in this important human pathogen (12).

## Materials and Methods

### Bacterial strains, media, and plasmids

*Shigella flexneri* strains were routinely grown on Trypticase soy agar (TSA; Trypticase soy broth [TSB] containing 1.5% [wt/vol] agar). When necessary, to ensure maintenance of the virulence plasmid, Congo red binding was examined on TSA plates containing 0.01% [wt/vol] Congo Red. Depending on the assay, liquid cultures of S. *flexneri* strains were routinely grown overnight at 30°C in either Luria-Bertani (LB) broth or minimal medium (M9 minimal medium supplemented with 0.4% D-glucose, 0.4% casamino acids, 0.01 mg/ml nicotinic acid, 0.01 mg/ml tryptophan, 0.01 mg/ml thiamine; 0.1 mM CaCl2; and 0.5 mM MgSO4) (46, 47). Overnight cultures were then diluted 1:100 and sub-cultured at 37°C with aeration in the specified medium (Note: 40 mM glycerol replaced D-glucose in minimal medium to allow induction of pBAD to occur; (48)). Where appropriate and throughout this study, diluted cultures were induced with 0.02% L-arabinose for the last 3 hours of a 5-hour growth period. These conditions were chosen because they allowed fusion proteins levels to mirror levels of native VirB at their peak in early stationary phase cultures, based on preliminary western blot analysis. To ensure plasmid maintenance, antibiotics were added to growth media at the following final concentrations: ampicillin, 100 μg/ml; chloramphenicol, 25 μg/ml. The bacterial strains and plasmids used in this study are listed in Table 1.

**Table 1.**
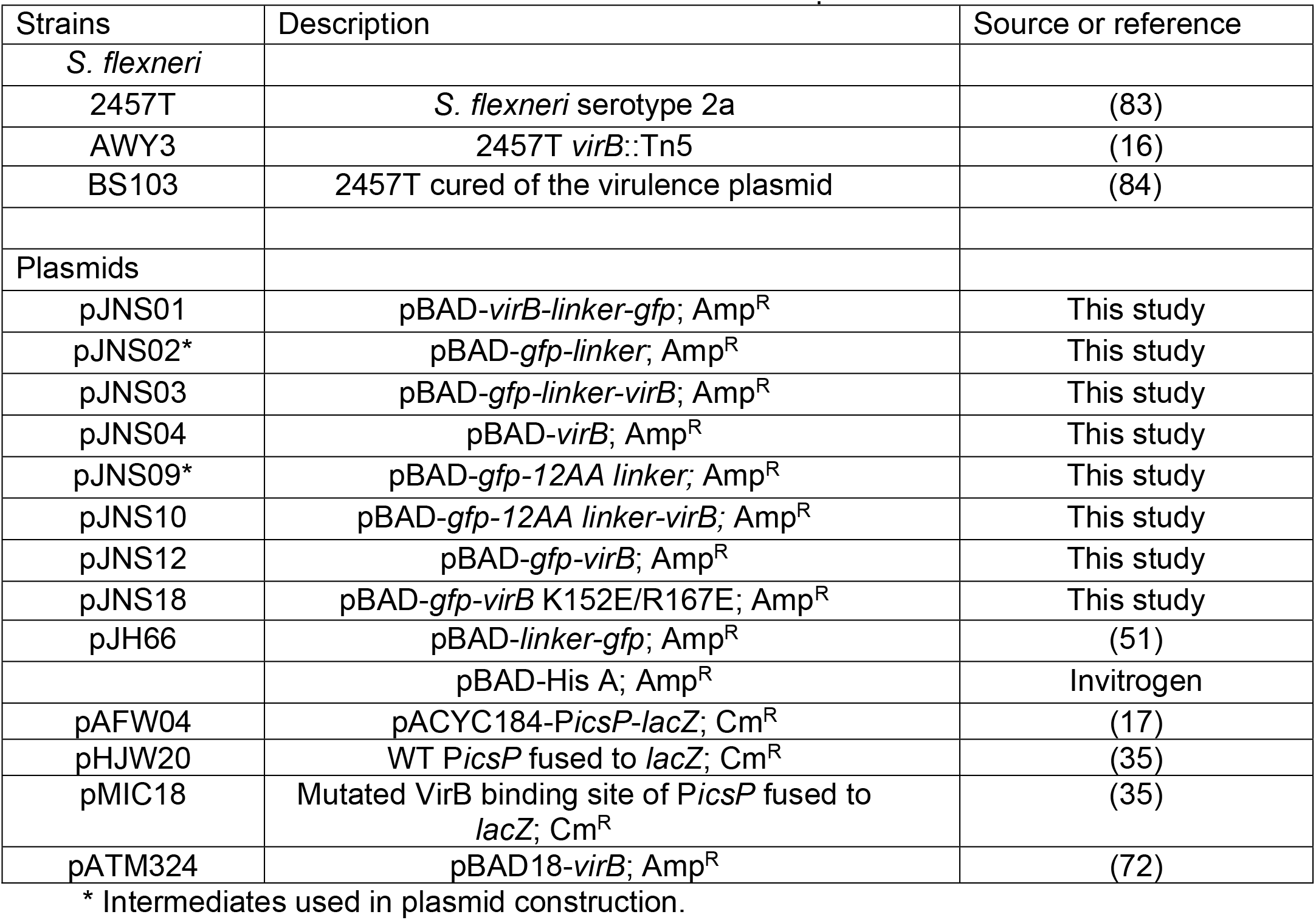
Bacterial strains and plasmids

| Strains | Description | Source or reference |
| --- | --- | --- |
| *S. flexneri* |  |  |
| 2457T | *S. flexneri* serotype 2a | (83) |
| AWY3 | 2457T *virB*::Tn5 | (16) |
| BS103 | 2457T cured of the virulence plasmid | (84) |
| Plasmids |  |  |
| pJNS01 | pBAD-*virB-linker-gfp*; Amp<sup>R</sup> | This study |
| pJNS02* | pBAD-*gfp-linker*; Amp<sup>R</sup> | This study |
| pJNS03 | pBAD-*gfp-linker-virB*; Amp<sup>R</sup> | This study |
| pJNS04 | pBAD-*virB*; Amp<sup>R</sup> | This study |
| pJNS09* | pBAD-*gfp-12AA linker*; Amp<sup>R</sup> | This study |
| pJNS10 | pBAD-*gfp-12AA linker-virB*; Amp<sup>R</sup> | This study |
| pJNS12 | pBAD-*gfp-virB*; Amp<sup>R</sup> | This study |
| pJNS18 | pBAD-*gfp-virB* K152E/R167E; Amp<sup>R</sup> | This study |
| pJH66 | pBAD-*linker-gfp*; Amp<sup>R</sup> | (51) |
|  | pBAD-His A; Amp<sup>R</sup> | Invitrogen |
| pAFW04 | pACYC184-*PicsP-lacZ*; Cm<sup>R</sup> | (17) |
| pHJW20 | WT *PicsP* fused to *lacZ*; Cm<sup>R</sup> | (35) |
| pMIC18 | Mutated *VirB* binding site of *PicsP* fused to *lacZ*; Cm<sup>R</sup> | (35) |
| pATM324 | pBAD18-*virB*; Amp<sup>R</sup> | (72) |

\* Intermediates used in plasmid construction.

### Plasmid constructs

The *gfp* used throughout this work encodes a monomeric superfolder GFP which contains key substitutions to minimize protein aggregation and enhance folding of both the fluorescent protein and the tagged protein of interest (49, 50). All pBAD inducible plasmids generated in this work were derived from pBAD/His A (Invitrogen). pJNS12 is *pBAD-gfp-linker-virB* flanked by NcoI and HindIII restriction sites. The *virB* gene in this plasmid, and others created in this work, was sourced from pATM324, while the *linker* and the *gfp* were identical to those found in pJH66 (51). During preliminary experiments (using pJNS03), a second active in-frame translation start site was identified upstream of the *virB* gene. This was removed from pJNS03 to generate pJNS12. pJNS18 is *pBAD-gfp-linker-virB* K152E/R167E and is derived from pJNS12. Each of the mutations encoding the substitutions are contained in a region flanked by BglII and PsiI sites in this construct. pJNS04 is *pBAD-virB,* where the *virB* gene is flanked by NheI and HindIII sites. pJNS01 is *pBAD-virB-linker-gfp* flanked by NcoI and HindIII restriction sites. Finally, pJNS10 is *pBAD-gfp-12AA-virB,* which is similar to pJNS12, however the 12AA linker (52) replaces the linker that is found in pJNS12. All plasmid constructs are listed in Table 1 and all were verified by Sanger dideoxy sequencing. The sequences of primers used in this study are available upon request.

### Analysis of fusion protein stability

Overnight cultures were diluted in LB and then induced, as described previously. Cells were normalized to cell density (OD600nm), harvested and washed with 0.2 M Tris buffer (pH 8.0), prior to resuspension in 200 μL 10 mM Tris (pH 7.4) and 50 μL 4X SDS-PAGE buffer (containing β-mercaptoethanol). Lysates were then sonicated to shear DNA and equal volumes of each normalized protein preparation were electrophoresed on 12.5% SDS-PAGE gels [Note: To preserve folding and fluorescence of proteins for the in-gel fluorescence assays, the samples were not boiled prior to electrophoresis.]. Due to the stability of the GFP even under denaturing conditions, in-gel fluorescence was detected using the Azure 500 (Azure Biosystems) using the Cy2 excitation setting of 492nm. For western blot analysis, GFP was detected using a polyclonal anti-GFP antibody obtained from Molecular Probes, and a GE anti-Rabbit IgG HRP (NA9340) secondary antibody, then imaged using chemiluminescence (Azure 500).

### Quantification of VirB fusion protein activity

For these assays, overnight cultures were diluted in LB and induced (described above), prior to being assayed. Culture were lysed and β-galactosidase activity was measured using a modified Miller protocol (16, 53).

### Visualization of fusion proteins & quantification of foci

To avoid growth medium autofluorescence, overnight cultures were diluted 1:100 in 2.5 ml minimal medium, induced as described previously and cells were immobilized in 1% agarose containing minimal media on glass slides before imaging. Data acquisition was performed using transmitted light and fluorescence microscopy (Zeiss Axio Imager M2) equipped with oil immersion (EC Plan-Neofluar 63X/Ph3). Fluorescence excitation was performed (X-cite 120LED) at a range of 470-525 nm. Images were captured by an ORCA Flash 4.0 LT Monochromatic Digital CMOS camera. Routinely, 5 fields of view were captured at random (solely using phase contrast during image focusing) for each experimental strain containing roughly 60 cells per field of view. Image analysis was completed using MicrobeJ (54). All cells in each field of view were counted and analyzed unless they did not fit the predetermined parameters (i.e. cell length > 4 μm or those that were too close to the border of field of view). Cell outlines were reviewed manually to ensure automatic segmentation was detected accurately. Also, focus detection was reviewed manually to determine whether signals were true foci or diffuse signals with maxima. Data was ordered by the number of foci or maxima detected followed by cell length, which was measured by the medial axis function in MicrobeJ. Cells greater than 4 μm in length likely had defects in cell division, and therefore, were categorized as “irregular” and removed from analyses.

## Results

### Characterization of VirB fused with GFP

To begin this study, VirB fused to msfGFP (monomeric, superfolder GFP (49, 50); referred to as GFP for the remainder of this paper) was expressed from the pBAD promoter found on the low copy (~15-20 copies per cell) plasmid, pBAD/His A (Table 1). Both N- and C-terminal VirB fusions were engineered and two different linkers were exploited (Fig. 1A) (52, 55). The plasmids generating VirB-GFP were abandoned in the early stages of this work, however, because protein production from one of these constructs did not positively correlate with L-arabinose induction (data not shown). Consequently, the two N-terminal fusions (GFP-VirB) were tested and compared to wild-type VirB for their ability to regulate a known, VirB-dependent promoter (16, 35). To do this, inducible plasmids pJNS04 (pBAD-VirB), pJNS10 (GFP-12AA-VirB), or pJNS12 (GFP-SGGGG-VirB), were introduced into a *virB* mutant derivative of *Shigella flexneri* carrying the *PicsP-lacZ* transcriptional reporter (17) and β-galactosidase activity was measured in cultures after induction. The data show that the GFP-VirB fusion with the SGGGG linker generated high *lacZ* expression similar to that observed with inducible VirB, suggesting that VirB remains functional in the context of the GFP fusion protein, whereas the ‘12AA linker’ appeared to interfere with the regulatory activity of VirB (Fig. 1B) (52). Consequently, GFP-SGGGG-VirB (encoded by pJNS12) was used for the remainder of this study.

**Fig. 1.**
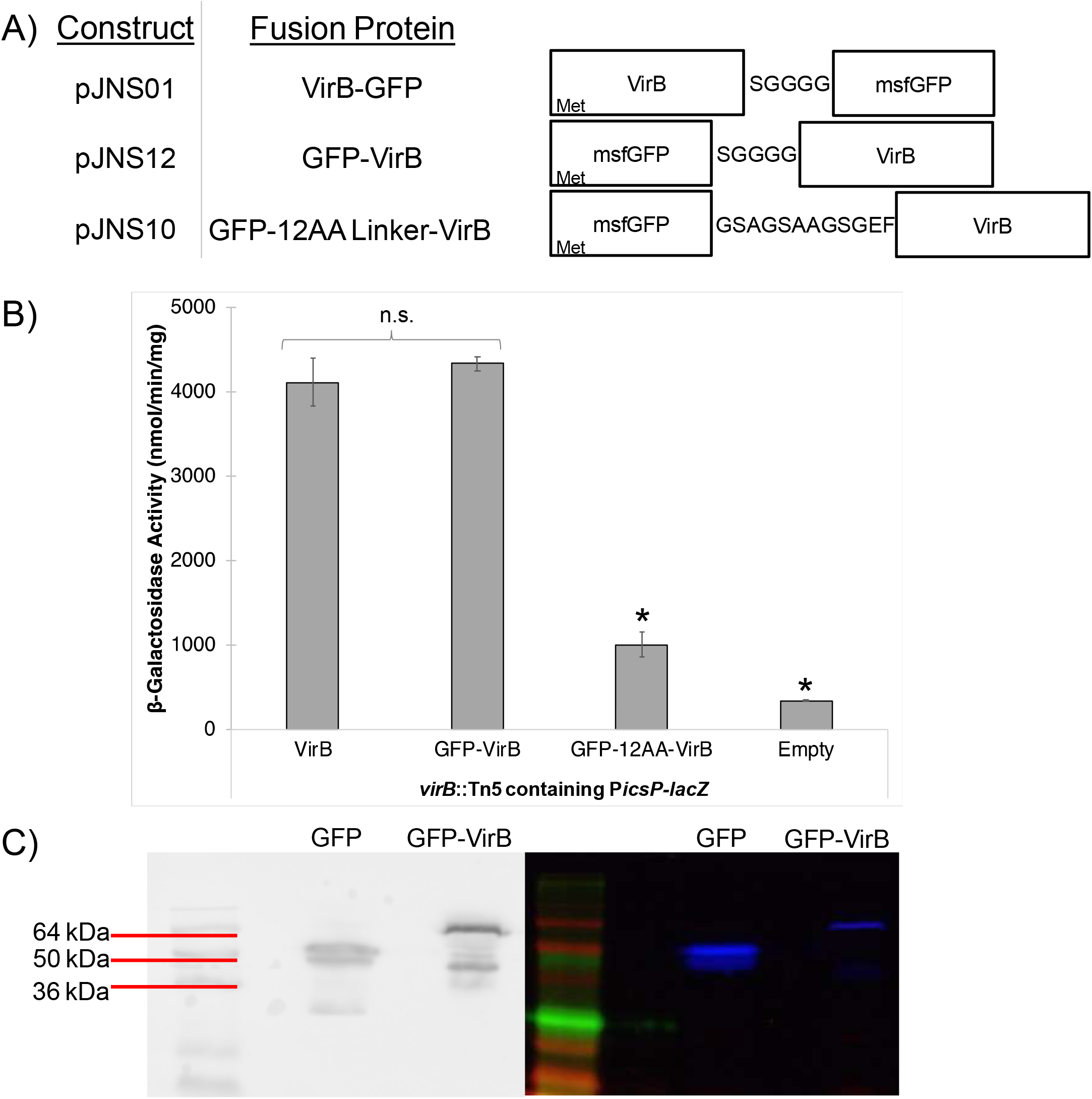
Constructs and proof of principle experiments to assess the activity and stability of VirB fusion proteins. (A) Constructs producing VirB fused to monomeric, superfolder GFP (msfGFP) at either the N-terminus or C-terminus using one of two linkers. (B) β-galactosidase assay used to assess the regulatory activity of GFP-VirB fusions with either a SGGGG (55) or “12AA” linker (52) at the VirB-dependent *icsP* promoter *(PicsP-lacZ;* pAFW04a). Student’s t-tests were used to measure statistical significance, * p < 0.05. β-Galactosidase activities generated in the presence of VirB (pATM324) and the GFP-VirB (pJNS12), were equivalent to those achieved with native VirB levels in previous work (35). (C) Western blot analysis using an anti-GFP antibody (left panel) and SDS-PAGE in-gel fluorescence (right panel) used to assess GFP-VirB protein stability. [Note: To preserve folding and fluorescence of proteins for the in-gel fluorescence assays, the samples were not boiled prior to electrophoresis. The potential for dimerization and formation of other higher order structures is elevated under these conditions].

To determine whether the observed regulatory activity of GFP-VirB at P*icsP-lacZ* could be attributed to the full-length fusion protein or a degradation product, protein production and stability of the GFP-VirB fusion protein was examined. To do this, two methods were used. First, western blots of whole cell lysates of cells producing GFP (pJH66) or GFP-VirB (pJNS12; grown using conditions identical to those used in the β- galactosidase assay) were probed using an anti-GFP antibody. The primary bands detected correspond to the GFP-VirB fusion protein (~62 kDa) and dimeric GFP with linker (~54 kDa), respectively (Fig. 1C – left panel). In each case more than 50% of the total protein detected was found in the uppermost band (full-length product), as determined by densitometry (data not shown). Second, to investigate the nature of the lower bands, and in the absence of a reliable VirB antibody, we chose to exploit the stability of full-length GFP under denaturing conditions. Using in gel fluorescence assays, fluorescent signals were detected from the upper bands, but not the lower bands (Fig. 1C – right panel), indicating that the lower bands were comprised of either truncated or truncated and misfolded protein. In sum, these analyses indicate that the fluorescent protein generated from pJNS12 primarily consists of the full-length GFP-VirB fusion and that the VirB moiety within this fusion is both stable and functional; a conclusion further supported by later experiments in this study (Fig. 5 & Fig. S1).

### GFP-VirB forms discrete foci in *Shigella* strains

The next step was to observe the fluorescent signal generated by the GFP-VirB fusion in S. *flexneri* cells bearing this construct. To do this, live cell imaging was utilized to avoid artifacts associated with cell fixing. First, we examined the GFP-VirB fusion protein, GFP, or negative control (empty plasmid control) in a S. *flexneri virB* mutant strain using identical induction conditions to those used in our previous proof of principle assays. Cells were live mounted and imaged using phase contrast and fluorescence microscopy.

As expected, cells carrying the empty plasmid control did not fluoresce (Fig. 2A bottom panel) and most cells producing GFP exhibited a single diffuse fluorescent signal, with a single distinct maximum [85% of all cells analyzed] (Fig. 2A middle panel; occasionally, two distinct maxima were seen, likely representing cells undergoing cell division [11% of analyzed cells]). Strikingly, when cells producing the GFP-VirB fusion were viewed, discrete fluorescent foci were observed in 98% of analyzed cells (Fig. 2A top panel). These foci were routinely seen at the quarter cell position (Fig. 2B). While the number of foci varied, commonly two, three, and four foci were detected during post-image analysis using MicrobeJ [30%, 37%, and 17%, respectively] (Fig. 2A). Notably, a weak positive correlation between the number of foci and cell length was found (R^2^ = 0.25, Fig. 2C), suggesting that the number of foci increase during cell elongation, prior to cell division.

**Fig 2.**
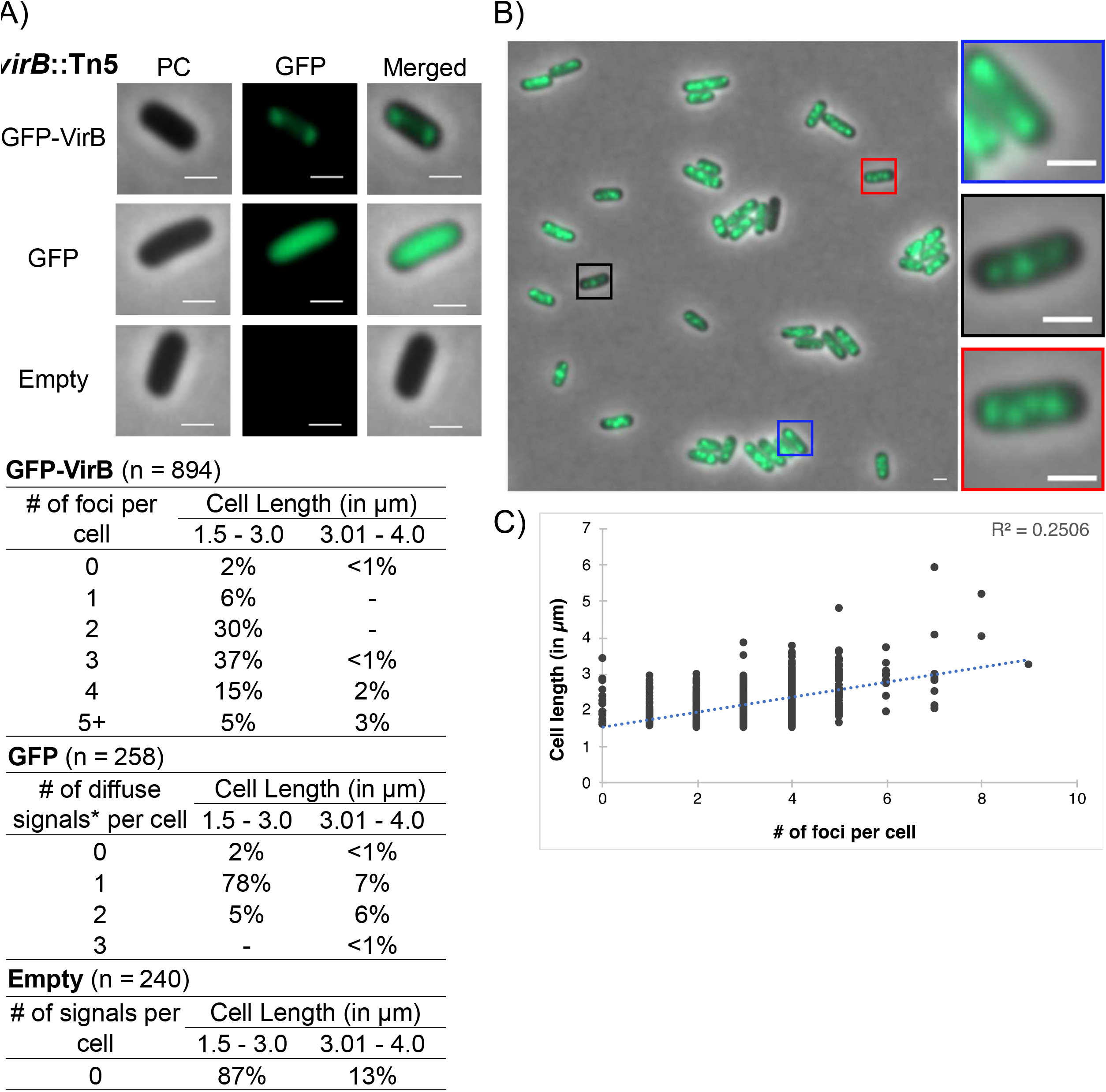
Live cell imaging of GFP-VirB in a *virB* mutant strain of S. *flexneri*. (A) Quantification of foci observed during live cell imaging of GFP-VirB in *virB* mutant S. *flexneri* using MicrobeJ. (-) indicates that the no cells were observed to fit the category in any of the images captured. Representative cells are shown. Phase contrast - PC (left column), fluorescence - GFP (middle column), and merged (right column) images of GFP-VirB (top row), GFP (middle row), and an empty plasmid control (bottom row). Scale bar represents 1 μm in all images. *Maxima detected by MicrobeJ. (B) A representative field of view. Magnified cell images show the number of foci commonly observed. Five fields of view were captured and analyzed across at least three independent replicates. (C) Scatterplot of cell length and number of foci observed in cells producing GFP-VirB.

To further investigate the subcellular location of VirB in the *Shigella* cytoplasm, we next chose to re-examine the GFP-VirB fusion in a wild-type S. *flexneri* background. In this strain background, data similar to those gathered in the *virB* mutant strain were obtained (99% of cells had 1 or more discrete focus/foci; longer cells had more foci than shorter cells; Fig. 3). This demonstrates that the presence of native VirB does not alter the localization pattern of GFP-VirB, but it also validates the localization pattern observed in the isogenic *virB* mutant strain of S. *flexneri.* Consequently, together these data (Fig. 2 & 3) provide novel insight into the VirB protein by finding that GFP-VirB is localized in the bacterial cell cytoplasm into 2-4 discrete foci at the quarter cell position. This raises the possibility that VirB forms previously undescribed hubs within in the bacterial cytoplasm, which may be critical for its role as a regulator of *Shigella* virulence.

**Fig. 3.**
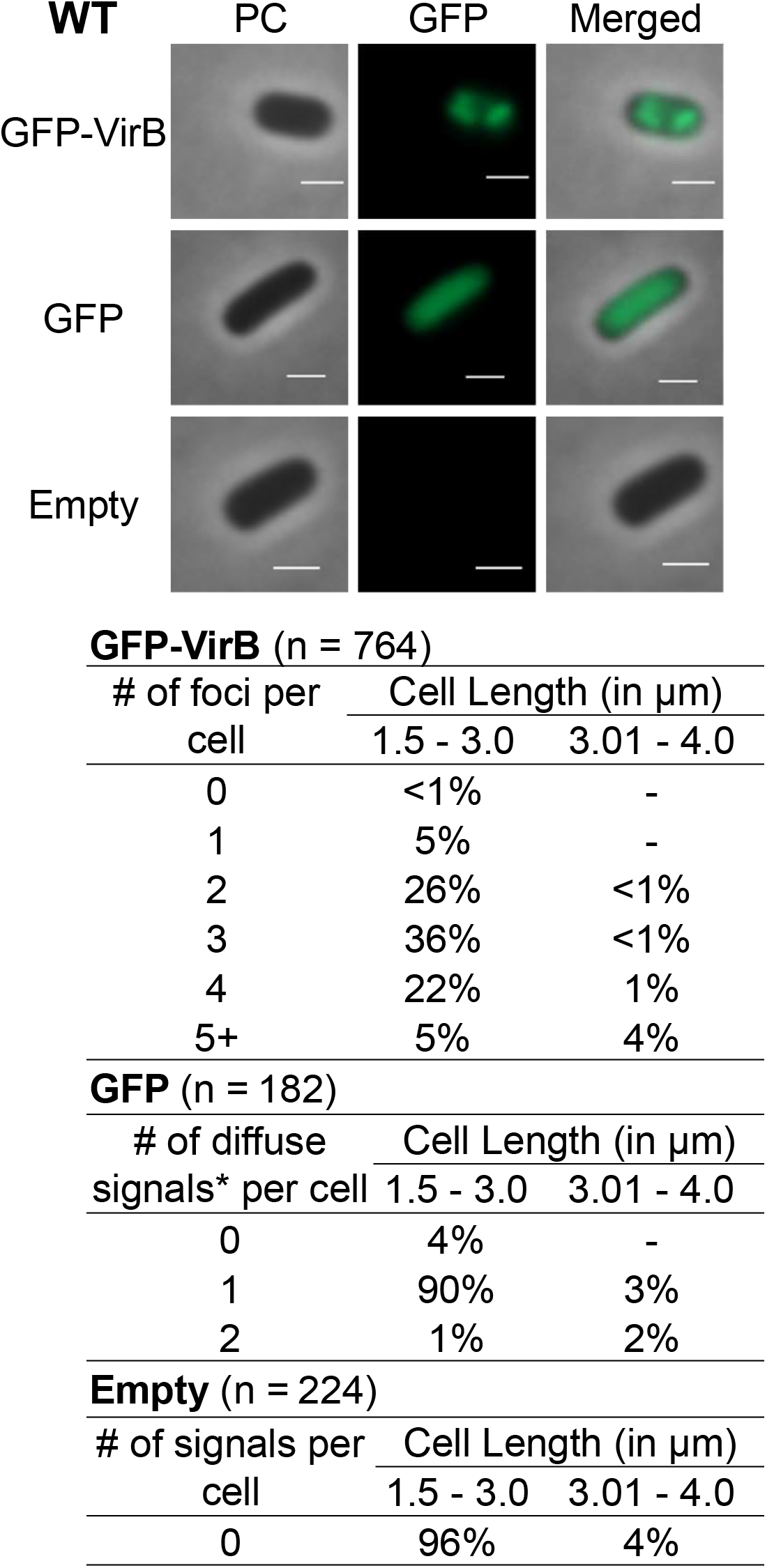
Live cell imaging of GFP-VirB in a wild-type strain of S. *flexneri*. Quantification of foci observed during live cell imaging of GFP-VirB in wild-type S. *flexneri* using MicrobeJ. (-) indicates that no cells were observed to fit the given category in any of the images captured. Representative cells are shown. Phase contrast (left column), fluorescence (middle column), and merged (right column) images of GFP- VirB (top row), GFP (middle row), and an empty plasmid control (bottom row). Scale bar represents 1 μm in all images. *Maxima detected by MicrobeJ.

### Focus formation of GFP-VirB is lost in the absence of pINV

Since VirB predominately regulates virulence genes carried by the large virulence plasmid pINV, we next chose to examine GFP-VirB focus formation in a strain lacking this large DNA molecule. To do this, we exploited a S. *flexneri* 2a derivative cured of pINV, BS103. In this strain background, GFP-VirB predominantly formed one or two diffuse fluorescent signals in the cytoplasm (Fig. 4, top panel; 24%, single maximum & 62%, 2 maxima). Critically, these signals did not resemble the tight foci observed in the *virB* mutant or wild-type strains (Fig. 2A & Fig. 3 top panel, respectively). Instead, the signals were much more diffuse, albeit less uniformly distributed than the GFP control (compare Fig. 4 - top panel to middle panel). Indeed, in this strain background the GFP-VirB fusion signal appeared to be nucleoid associated rather than distributed evenly through the cytoplasm (Fig. 4 - top panel; evident in merged image). Again, control experiments yielded expected results in these assays (Fig. 4, middle & bottom panel).

**Fig. 4.**
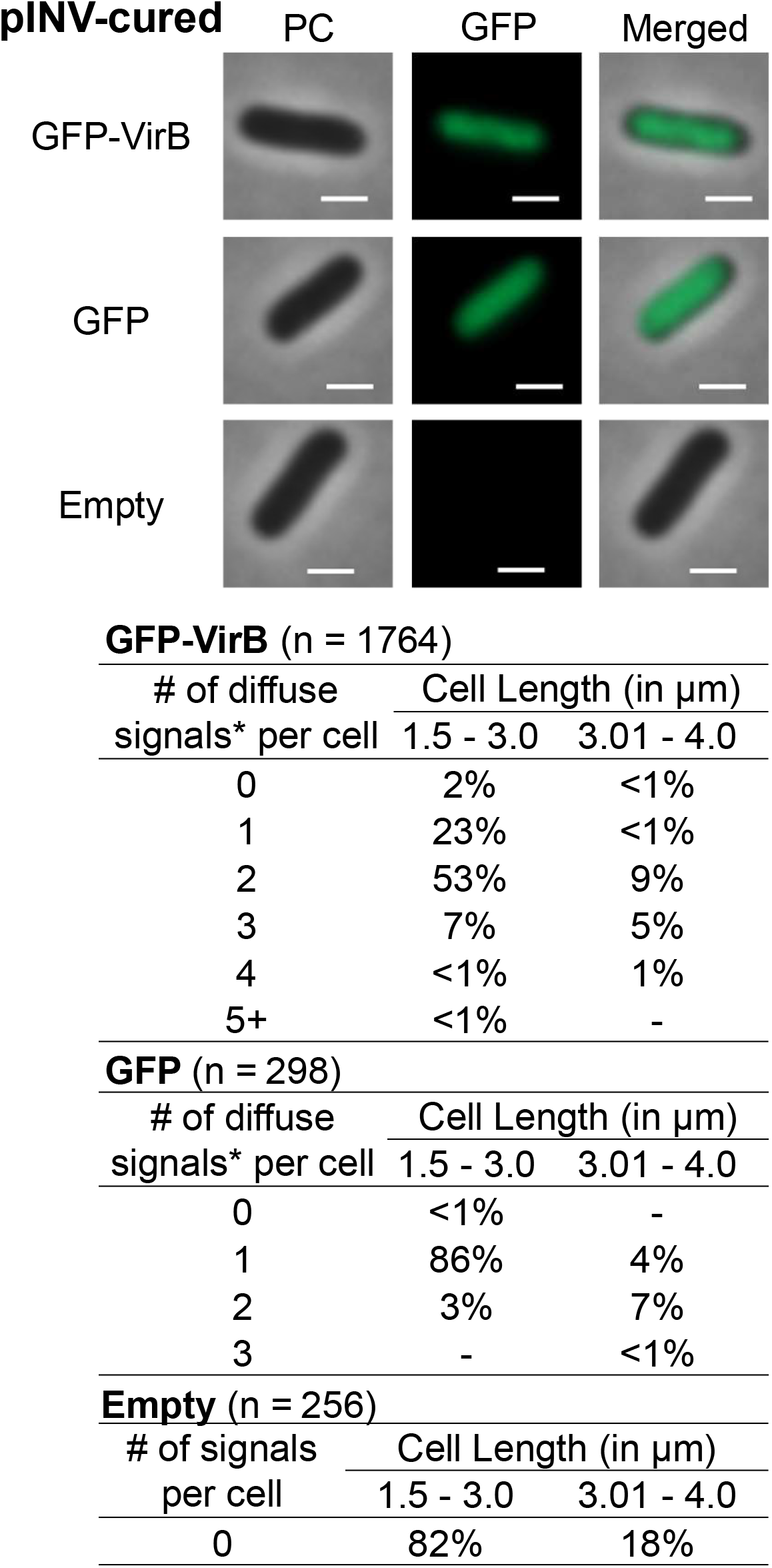
Live cell imaging of GFP-VirB in a pINV-cured strain of S. *flexneri*. Quantification of fluorescent signals observed during live cell imaging of GFP-VirB in pINV-cured S. *flexneri* using MicrobeJ. Note, only diffuse signals were observed. *Maxima detected by MicrobeJ. (-) indicates that that no cells were observed to fit the given category in any of the images captured. Representative cells are shown. Phase contrast (left column), fluorescence (middle column), and merged (right column) images of GFP-VirB (top row), GFP (middle row), and an empty plasmid control (bottom row). Scale bar represents 1 μm in all images.

### A DNA-binding mutant derivative of GFP-VirB does not form foci in S. *flexneri*

The finding that GFP-VirB did not form discrete foci in the absence of pINV raised the possibility that direct VirB interactions with pINV were required for focus formation in wild-type and *virB* mutant strain backgrounds. To test this hypothesis, the fluorescence of a GFP-VirB fusion carrying amino acid substitutions in the DNA binding surface of VirB was examined. To do this, two amino acid substitutions, K152E and R167E (56, 57), were introduced into the helix-turn-helix DNA binding domain of VirB in the fusion (Fig. 5A). Each of these substitutions had previously been shown to prevent VirB from binding to DNA and, hence, prevent regulation by VirB (56, 57). The stability of the resultant fusion protein, GFP-VirB K152E/R167E, and its ability to regulate transcription was then examined (Fig. S1A). Again, little protein degradation was observed by western blot analysis and in-gel fluorescence, suggesting that GFP-VirB K152E/R167E was as stable as GFP-VirB (Compare Fig. 1C to Fig. S1A). But, as predicted by earlier studies (56, 57), this protein was unable to regulate P *icsP-lacZ.* Indeed, levels of β-galactosidase activity in the presence of GFP-VirB K152E/R167E were comparable to those obtained with the empty control (Fig. S1B). The finding that two amino acid substitutions in the DNA binding domain of the VirB moiety are sufficient to destroy the regulatory activity of this fusion (Fig. S1B) is not only consistent with previous findings (56, 57), but further supports our assertion that the VirB moiety remains properly folded in the context of the fusions used in this study.

**Fig. 5.**
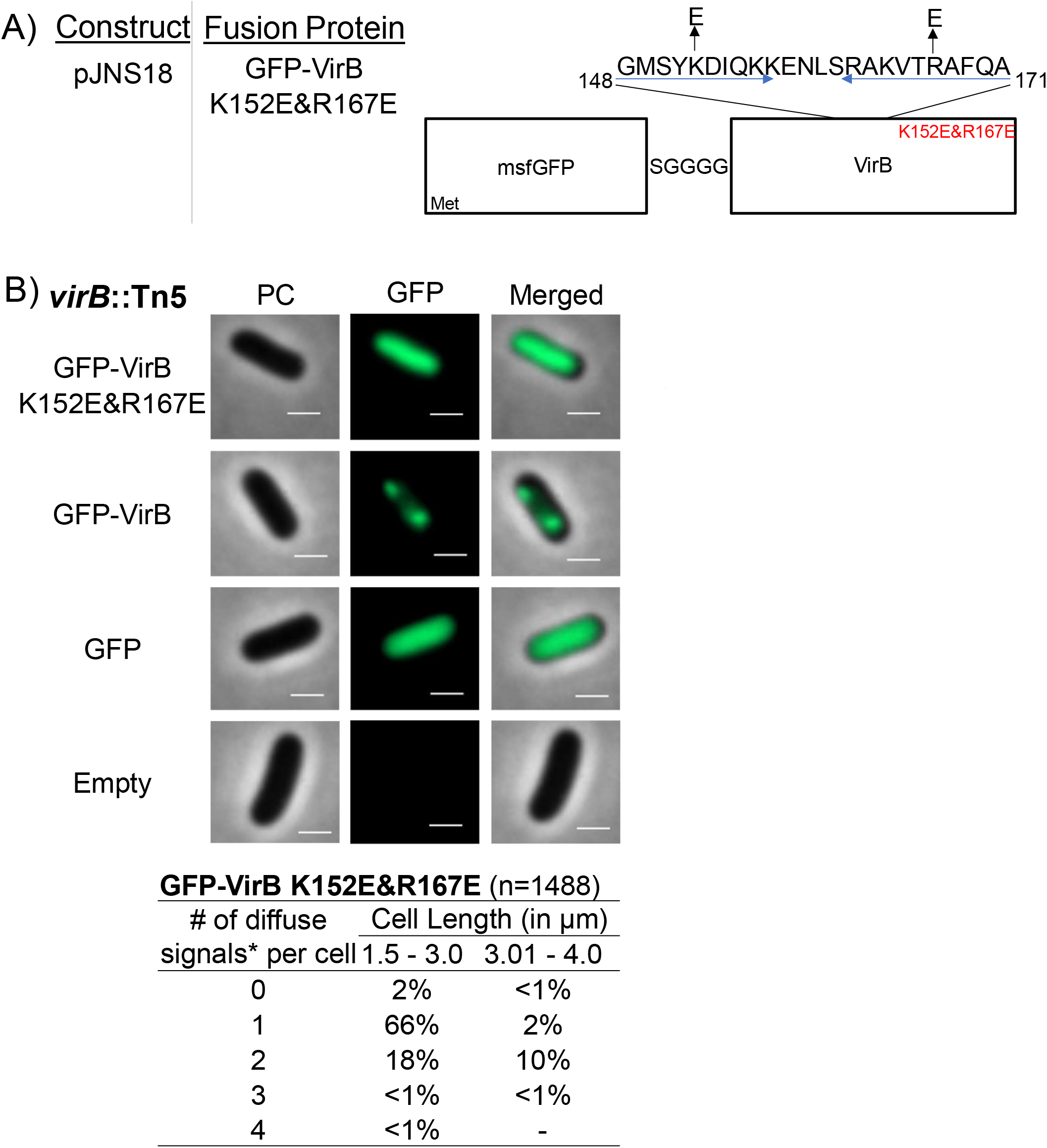
Construction of a GFP-VirB DNA-binding mutant and live cell imaging of this fusion in a *virB* mutant strain of S. *flexneri*. (A) Construct producing GFP-VirB with two amino acid substitutions K152E and R167E in the helix-turn-helix DNA-binding domain (denoted as blue arrows; adapted from (56)) (B) Quantification of fluorescent signals observed during live cell imaging of GFP-VirBK152E/R167E in *virB* mutant S. *flexneri* using MicrobeJ. Note, diffuse signals observed for this fusion were detected as maxima by MicrobeJ. (-) indicates that that no cells were observed to fit the given category in any of the images captured. Representative cells are shown. Phase contrast (left column), fluorescence (middle column), and merged (right column). Scale bar represents 1 μm in all images.

When the GFP-VirB KE152/R167E fusion was examined in the *virB* mutant, wild-type, and pINV-cured strain using fluorescence microscopy, a diffuse fluorescent signal was observed (Fig. 5B – top panel & Fig. S1C). This signal was evenly distributed throughout the cytoplasm and was indistinguishable from the signal emitted from cells producing GFP (Fig. 5B - compare top panel to 3^rd^ panel). Importantly, no discrete foci were observed with the GFP-VirB KE152/R167E expressing construct. Consequently, we concluded that an active DNA-binding domain of VirB is required for VirB to form foci in the cell cytoplasm. Moreover, in combination with the findings made with GFP-VirB in a pINV-cured strain, these outcomes strongly suggest that VirB:DNA interactions with pINV are responsible for the focus formation observed in the wild-type and *virB* mutant derivative of S. *flexneri;* that these foci are not a caused by GFP aggregation.

### A small plasmid carrying a VirB binding site restores focus formation in pINV-cured *Shigella*

Finally, because focus formation was dependent on the presence of pINV, we next chose to examine if a small surrogate plasmid carrying a VirB binding site would allow fluorescent foci to form in a pINV-cured background of S. *flexneri.* To test this, a low copy plasmid, pHJW20 (35) carrying a well-characterized VirB binding site; (22, 35, 36)) or a derivative pMIC18 (35), with a mutated VirB binding site was introduced into BS103 and then either pJNS12, pJNS18, or one of the control plasmids (pJH66 or pBAD/His A) were introduced.

Remarkably, foci formed in the presence of the small surrogate plasmid carrying the wild-type VirB-binding site (Fig. 6A) but not with the derivative carrying the mutated site (Fig. 6E). Furthermore, the DNA-binding mutant derivative remained diffuse regardless of whether a wild-type or mutated site was present (Fig. 6F), thus strengthening support for the previous interpretation that the ability of VirB to interact with DNA is absolutely necessary in order for foci to form (Fig. 6B). Once again, the other included controls behaved as expected (Fig. 6B-D & 6F-H).

**Fig. 6.**
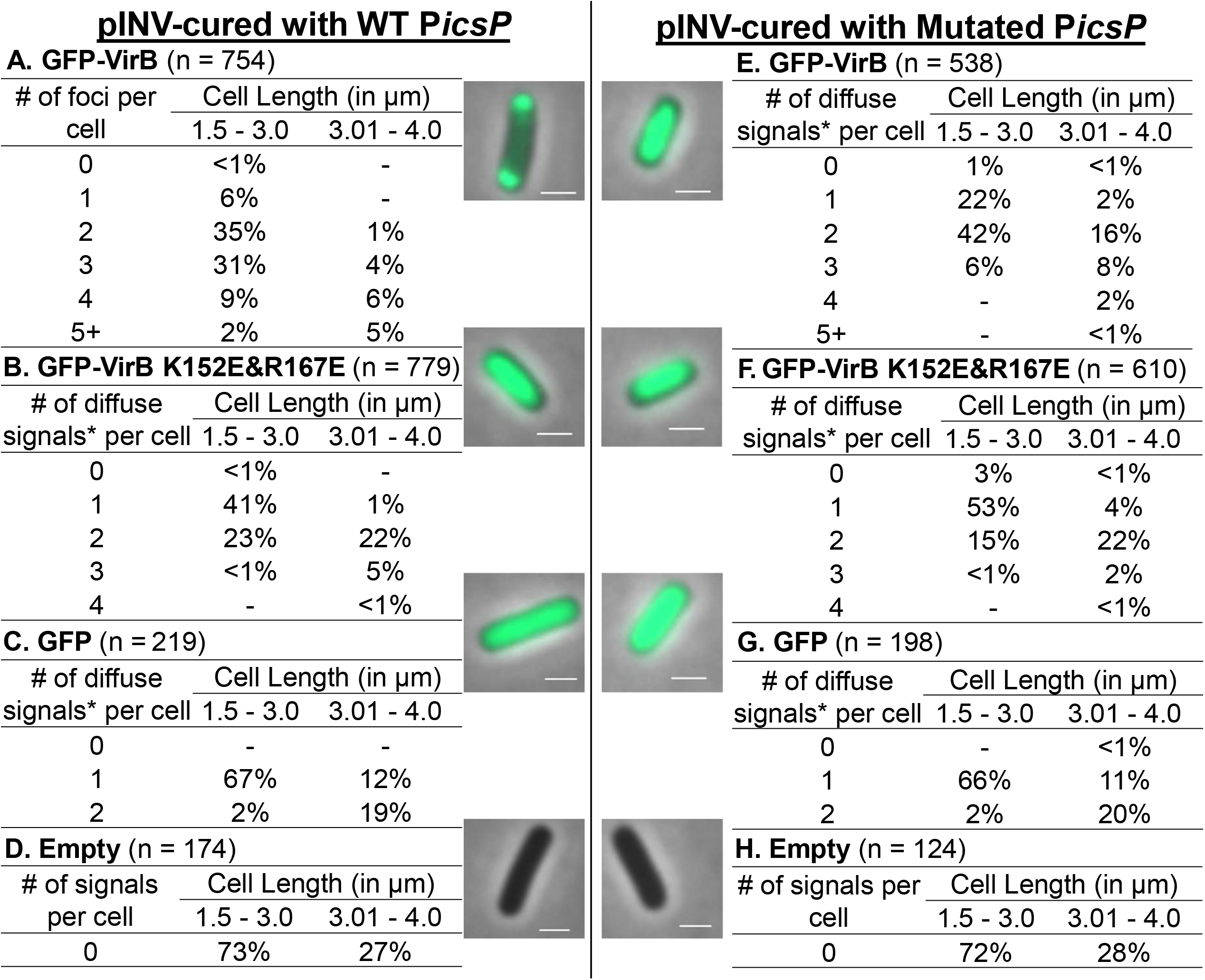
Live cell imaging of GFP-VirB and controls in a pINV-cured strain of S. *flexneri* carrying a plasmid bearing a wild-type or mutated VirB-binding site. Quantification of fluorescent foci of GFP-VirB (A) or *maxima (B, C, E-G) associated with diffuse signals observed during live cell imaging using MicrobeJ. No fluorescent signal was detected for the empty controls (D & H). (-) indicates that no cells were observed to fit the given category in any of the images captured. Representative merged cell images of pINV-cured S. *flexneri* cells containing pHJW20 (WT *PicsP)* (AD) or pMIC18 (Mutated *PicsP)* (E-H) are shown. Scale bar represents 1 μm in all images.

Our findings demonstrate that interactions between VirB and its cognate DNA binding sequence are necessary and sufficient for the formation of discrete cytoplasmic GFP-VirB foci in S. *flexneri.* Further, the results show GFP-VirB can form foci independently from any pINV-encoded factors, other than the native VirB-binding sites found on this molecule. In addition to providing new information on VirB activity, these observations raise the possibility that this fluorescent fusion protein may be used in conjunction with its cognate binding sequence as a plasmid or genome marker for fluorescent microscopy experiments in *Enterobacteriaceae* and other species.

## Discussion

In this study, we have investigated the subcellular location of VirB, a key transcriptional regulator of *Shigella* virulence plasmid genes (13, 19), using a GFP-VirB fusion protein (Fig. 1). Our data show that GFP-VirB forms discrete foci in the *Shigella* cytoplasm (Fig. 2 & 3). Three lines of evidence demonstrate that these foci are completely dependent upon VirB engaging its DNA binding site; i) the loss of GFP-VirB foci in pINV-cured cells (Fig. 4), ii) the loss of foci when a DNA-binding mutant derivative of GFP-VirB was produced in a *virB* mutant derivative (Fig. 5), and iii) the restoration of GFP-VirB foci when a small surrogate plasmid bearing a VirB recognition site was introduced into pINV-cured cells (Fig. 6). Throughout this study, GFP-VirB levels were matched to native levels of VirB (Materials and Methods; data not shown) and the GFP-VirB fusions were found to remain stable and active under the conditions of our assays (Fig. 1C & B, respectively). Thus, our work strongly suggests that VirB forms functional hubs on DNA after engaging its recognition site, and this occurs in the absence of any other pINV-encoded factor. The reason for hub formation, the macromolecular nature of these hubs, and the role that this plays in transcriptional anti-silencing of virulence genes on pINV are discussed below.

The regulatory actions of VirB are thought to be limited to the *Shigella* virulence plasmid (13, 15, 58). Consequently, we were surprised to find that VirB formed foci in the bacterial cytoplasm for several reasons. Firstly, the *Shigella* virulence plasmid, pINV, is large (230 kb) (59, 60), approximately 1/20^th^ of the *Shigella* chromosome. Secondly, VirB up-regulates at least 50 virulence-associated genes that are scattered around pINV (16, 61, 62) and found within the large *ipa-mxi-spa* operons (15, 22, 63, 64). Thirdly, another transcription factor that binds to pINV, VirF, does not form foci in *Shigella* when fused to mCherry (data not shown). Although transcriptional regulators have been severely understudied in terms of their localization, especially in prokaryotic systems (37), the paradigm has always been that these proteins are found throughout the bacterial nucleoid at concentrations too low to be detected by standard fluorescence microscopy. Consequently, our finding that the GFP-VirB was forming DNA-dependent fluorescent foci in the bacterial cytoplasm was intriguing because it suggested that VirB was forming multimeric complexes on DNA.

So, why then would a transcriptional regulator of virulence genes in *Shigella* form multimeric complexes when it engages DNA? A clue came from the evolutionary lineage of VirB. VirB belongs to the ParB superfamily (19, 56); a family of proteins where the vast majority play central roles in plasmid or chromosome segregation. Examples of this family have been studied in depth and are widespread among well-studied bacterial phyla (38, 42, 45). Some shared features of VirB and ParB are a centrally-located, highly-conserved, helix-turn-helix DNA binding domain, and a C-terminally-located dimerization domain (56, 65). Additionally, both proteins recognize and bind to similar, but non-identical sites, which contain a 14 bp inverted repeat (22, 35, 36, 66).

ParB and other family members, including Spo0J, KorB, SopB, and RepB, display a discrete subcellular localization in the bacterial cytoplasm in experiments similar to those described here. Frequently, 2, 3 or 4 foci are routinely formed by ParB fusions and, like VirB, the precise number appears to correlate with cell length (41). Initially, ParB was thought to engage its *parS* site and then, spread along DNA to form a DNA:ParB filament (43, 67, 68). More recently, however, the assembly of ParB at its *parS* recognition sites has been investigated using single molecule approaches (45) and super-resolution microscopy (69). These studies reveal that ParB:ParB interactions, described as ParB bridges, are commonplace on DNA (45) and that *in vivo* almost all ParB molecules are actively confined around *parS* sites by a network of synergistic protein-protein and protein-DNA interactions (69). These data indicate that rather than binding only to specific sequences and subsequently spreading along DNA (43, 67, 68), ParB seems to form stochastic DNA cages when ParB proteins interact from distal *parS* sites (69). Given the relatedness of VirB to ParB, the dynamic multimeric ParB complexes and ParB:DNA lattices reported by Sanchez *et al.,* and others are intriguing, because they raise the possibility that similar VirB complexes may be responsible for the GFP-VirB foci that we observe in this study. At this stage, it remains unclear if VirB spreads along DNA once bound to its recognition sites (36) or if it forms multimeric complexes with itself once bound to DNA (akin to the ParB-mediated bridging or caging described in (45, 69)). However, the number of fluorescent foci per cell observed in our assays is far fewer than the estimated number of VirB-regulated loci on pINV, and so we currently favor the VirB bridging/caging hypothesis.

Our studies of VirB-dependent transcriptional regulation in *Shigella* have already revealed that VirB is not a traditional transcription factor (23). VirB can regulate transcription from remotely located binding sites >1kb upstream of a promoter; (18, 23, 35) and instead of promoting the recruitment or activity of RNA polymerase, like other transcription factors, VirB functions to offset or counteract transcriptional silencing imparted by the pervasive nucleoid structuring protein, H-NS (15, 16, 22, 23). The parallels between VirB and ParB revealed by our work are tantalizing and may provide new insight into the transcriptional anti-silencing mechanism imparted by VirB. For instance, multivalent self-interactions among VirB molecules could bring multiple VirB-binding sites together, resulting in clusters of assembled segments with extruding loops of DNA. In some ways, this notion resembles recent discoveries of transcription factor complexes in eukaryotic nuclei (70) and means that VirB may be functioning as a domain organizing protein (71), but one that is apparently designated for pINV. Such large scale molecular interactions between VirB molecules docked on pINV may lead to the remodeling of the AT-rich DNA molecule pINV, leading to the eviction or remodeling of H-NS:DNA interactions such that gene expression occurs. Ongoing research in our laboratory is exploring these hypotheses.

In addition, the relatedness of VirB to ParB also raises questions about the potential for VirB to function in plasmid partitioning. Although VirB is a close homologue of the ParB encoded by S. *flexneri* pINV, ParBSf (37% identical) there is currently no evidence that these proteins functionally substitute for one another: in cells lacking *virB,* virulence gene expression does not occur (13, 15) (even though ParBSf is encoded by pINV) and the plasmid pINV is stably maintained (72). Additionally, the VirB protein does not contain the N-terminal domain found in ParB that is required for interaction with the ATPase partner protein, ParA, (22, 65) and no *parA* gene is found in the vicinity of the *virB* gene or elsewhere on pINV (with the exception of *parA* – encoding the ParASf that works with the designated ParBSf). Instead, all evidence indicates that the role of VirB is to modulate gene expression by antagonizing H-NS mediated silencing (22, 73). Nevertheless, based on findings presented in this study, the interplay between VirB and the ParABSf system seems worthy of further exploration. For instance, it would be interesting to determine if VirB and ParB co-localize in S. *flexneri* cytoplasm and if there is any evidence of molecular crosstalk between VirB and the native ParAB system that leads to VirB-mediated interference in pINV partitioning. These experiments may help to elucidate why overexpression of VirB leads to the destabilization of pINV (72). While the instability of pINV was initially proposed to be caused by the additional VirB-dependent transcriptional burden placed on *Shigella* cells (74), our new data raise the possibility that VirB might directly interfere with the ParAB system on pINV. Since recent work has revealed that the pINV toxin:antitoxin systems are the dominant means by which pINV is maintained in *Shigella* cells (75, 76), if VirB is found to interfere with the ParAB system, this may have necessitated this unusual reliance on the pINV-encoded toxin:antitoxin systems.

Our finding that GFP-VirB foci still form in *Shigella* when a small surrogate plasmid bearing a single VirB recognition site was introduced into pINV-cured cells, demonstrated that focus formation was not reliant on anything found on pINV, except the VirB binding sites. At this stage, it remains unclear why foci were seen to form on these small pACYC plasmids. These foci could be caused by VirB:VirB bridging interactions across different copies of pACYC184 (a medium copy plasmid), the clustering of medium-copy plasmids that has been previously reported (77, 78), or some combination thereof. Regardless, it is clear from our work that the DNA binding activity of VirB is for these foci to appear. These findings are interesting for another reason; they raise the possibility that GFP-VirB and its recognition site could be used in combination as markers of DNA molecules, thus expanding the repertoire of ParB/parS-like markers that have been used in this manner (79). Indeed, unique features of the VirB/binding site system may avoid some of the lethal segregation defects that can result when *parS* sites are built into bacterial chromosomes at a distance (80). Since the VirB recognition site is only 25 bp long, the resulting system would allow for a drastic reduction in the length of foreign DNA that is required to be introduced into chromosomal or plasmid DNA of interest. Notably, a tandem array of over 250 *lacO* sites has often been used to generate a visible focus of fluorescently tagged LacI on bacterial genomes (78, 81,82). Future work will determine if a single 25 bp VirB binding site taken out of its context in the *icsP* promoter sequence, and placed onto a pACYC plasmid is sufficient to allow focus formation (the current plasmid contains ~1kb worth of *icsP* promoter sequence), and if this potential marker of DNA works reliably in closely related enteric organisms and other more distantly related bacteria.

In sum, this study provides a new view of a major regulator of virulence gene expression in *Shigella* spp. The formation of GFP-VirB foci strongly suggests that VirB forms large VirB:VirB and VirB:DNA complexes once it engages its DNA recognition sites, raising the possibility that VirB functions as a major remodeler of pINV topology. This is tantalizing given that the established role of VirB is to counteract transcriptional silencing imparted by another DNA structuring protein, H-NS. In addition, our work introduces the idea that fluorescently tagged VirB and its binding site may have utility as a DNA marker, allowing plasmid or perhaps even chromosomal loci to be tracked in cells. Finally, this work also fosters questions about the evolution of VirB to its role as a transcriptional anti-silencer from a DNA partitioning protein. Consequently, this study ties together the fields of virulence gene regulation, plasmid partitioning, and possibly even plasmid topological remodeling in bacterial cells. We anticipate that the work presented here will be foundational for future studies that explore these relationships and activities in more detail, and think this study, once again, highlights the merits of taking a holistic view of molecular biology by examining the subcellular localization of proteins, like VirB, in whole, living cells.

## Acknowledgements

We thank Monika M. Karney and Dr. Boo Shan Tseng for technical and microscope support. We are grateful to members of the Wing lab past and present for many helpful discussions on this research topic. The University of Nevada Las Vegas (UNLV) Genomics Core Facility (sponsored by the National Institutes of General Medical Sciences; P20GM103440) provided sequencing services. This work was supported by the National Institute of Allergy and Infectious Diseases at the National Institutes of Health (NIH), R15 AI090573 (H.J.W) and by the National Institute of General Medical Sciences at the National Institutes of Health (NIH) R01GM118792-01 (G.B.).The content is solely the responsibility of the authors and does not necessarily represent the official views of NIH. J.N.S. was the recipient of a GPSA travel grant that supported travel to the University of Wyoming for image capture training appropriate for this study. The authors declare no conflicts of interest.

---

**Supplementary Fig. 1.**
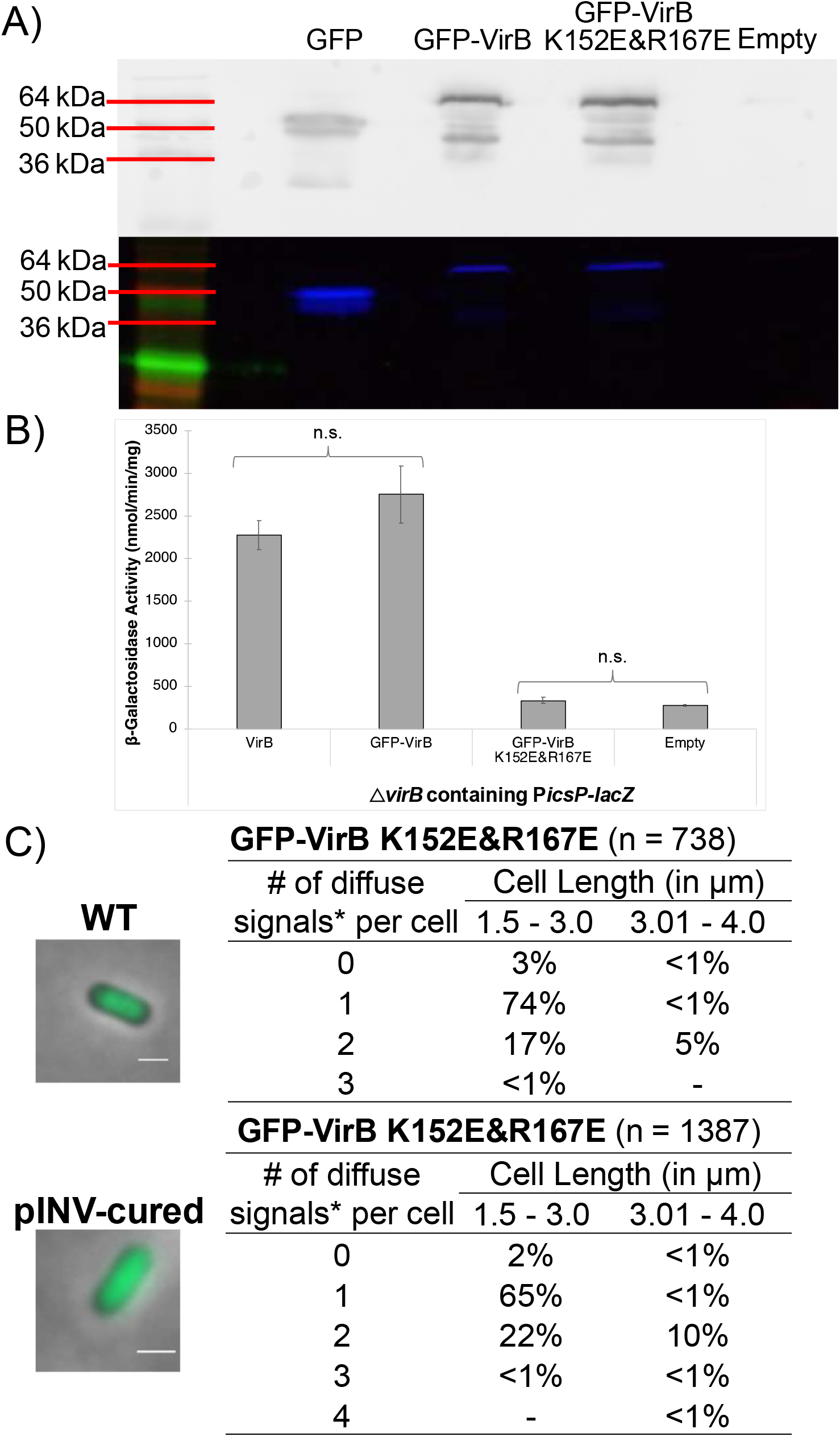
Proof of principle experiments to assess the activity and stability of GFP-VirB K152E/R167E & live cell imaging in wild-type and a pINV-cured strain of S. *flexneri.* (A) Western blot using an anti-GFP antibody (top panel) and SDS-PAGE in-gel fluorescence (bottom panel) used to assess fusion protein stability. [Note: To preserve folding and fluorescence of proteins for the in-gel fluorescence assays, the samples were not boiled prior to electrophoresis. The potential for dimerization and formation of other higher order structures is elevated under these conditions]. (B) β-galactosidase assay used to compare the regulatory activity of GFP-VirB fusions at the VirB-dependent *icsP* promoter (*PicsP-lacZ;* pAFW04a). Student’s *t*-tests were used to measure statistical significance, * p < 0.05. (C) Live cell imaging and quantification of diffuse fluorescence signal generated by GFP-VirB K152E/R167E in wild-type and pINV-cured S. *flexneri.* (-) indicates that that no cells were observed to fit the given category in any of the images captured. Scale bar represents 1 μm in all images. *Maxima detected by MicrobeJ.

